# Energetic decoupling of phycobilisomes from photosystem II involved in nonphotochemical quenching in red algae

**DOI:** 10.1101/2022.01.21.477255

**Authors:** Yu-Hao Chiang, Yu-Jia Huang, Han-Yi Fu

## Abstract

To mitigate photodamage under fluctuating light conditions, photosynthetic organisms respond by regulating light energy absorbed by light-harvesting complexes and used for photochemistry. Nonphotochemical quenching acts as a frontline response to prevent excitation energy from reaching the photochemical reaction center of photosystem II. The mechanisms underlying nonphotochemical quenching in red algae, which display unique combination of light-harvesting transmembrane antenna proteins and membrane-attached phycobilisomes, appear to be different from those in cyanobacteria, green algae, and plants. Several single-process models have been proposed for red algal nonphotochemical quenching, yet the possibility of more than one process being involved in nonphotochemical quenching awaits further investigation. To assess multiple nonphotochemical quenching processes in the extremophilic red alga *Cyanidioschyzon merolae*, fluorescence analyses with light preferentially absorbed by phycobilisomes or photosystems were utilized. Energetic decoupling of phycobilisomes from photosystem II and intrinsic photosystem II quenching were identified as two dominant processes involved in nonphotochemical quenching and distinguished by their kinetics. Whereas the degrees of energetic decoupling remained similar after its induction, the degrees of intrinsic photosystem II quenching varied depending on the illumination period and intensity. The respective effects of protein crosslinkers, osmolytes, ionophores, and photosynthetic inhibitors on the kinetics of nonphotochemical quenching suggested that the energetic decoupling involved conformational changes associated with the connection between the PBS and PSII. Furthermore, the surface charge on the thylakoid membrane played a significant role in the modulation of red algal nonphotochemical quenching.

**One-sentence summary:** Energetic decoupling of phycobilisomes from photosystem II and intrinsic photosystem II quenching were involved in nonphotochemical quenching of the extremophilic red alga Cyanidioschyzon merolae.

## INTRODUCTION

Effective utilization of light energy for photochemical reactions under variable environments with different light qualities and quantities is crucial in photosynthesis. The capacity of light energy reaching a photochemical reaction center, also called the light-harvesting antenna size, is enhanced by the supplement of light-absorbing antenna pigments excitonically coupled to the reaction center pigments. These antenna pigments are located within photosystems and peripheral light-harvesting antenna proteins. Whereas the antenna pigments facilitate photochemical reaction, the reaction center faces the risk of photodamage when the excitonic flux toward the reaction center exceeds the capacity of photochemistry. Such a high excitonic flux toward the reaction center results in the accumulation of reactive oxygen species and ultimately reduces photosynthetic capacity. PSII is more sensitive to light than PSI in most of the stress conditions, and several protective responses have been identified as measures to counteract PSII photodamage at different stages (Sonoike, 2011; Derks et al., 2015). To protect PSII from overexcitation, nonphotochemical fluorescence quenching (NPQ) serves as a frontline response and its level reflects the extent of the overall nonradiative deactivation process of excitons reaching the reaction center.

Extensive research on plants, green algae, and cyanobacteria has revealed diverse NPQ mechanisms related to the constituents of the photosynthetic apparatus. Vascular plants and green algae possess transmembrane light-harvest complexes (LHCs) as light-harvesting antennas and exhibit the process qE, a major component of NPQ defined by its induction driven by a transmembrane proton gradient of thylakoids. Whereas induction of qE requires the PsbS protein in vascular plants (Li et al., 2000), the protein essential for green algal qE is the LHCSR protein rather than PsbS (Peers et al., 2009; Tibiletti et al., 2016). The variety of mechanisms underlying qE in different photosynthetic organisms was also exemplified by how the quenching sites are located differently: within both LHCII and PSII of vascular plants but within only LHCII of green algae (Nicol et al., 2019; Tian et al., 2019). Cyanobacteria employ membrane-attached phycobilisomes (PBSs) instead of LHCs for light harvesting. They lack qE but possess blue light-driven NPQ involving activated orange carotenoid proteins binding to the PBS core (Wilson et al., 2006). Another component of NPQ, qT, is related to the state transitions during which the exitonic energy between PSII and PSI are redistributed. In vascular plants and green algae, the induction of qT refers to LHCIIs detaching from PSII and moving to PSI when PSII is preferentially excited, and the process is reversed when PSI is preferentially excited (Ruban and Johnson, 2009). The lateral migration of LHCIIs toward PSI or PSII is determined by respective phosphorylation or de-phosphorylation of LHCIIs, and the kinase responsible for phosphorylation of LHCIIs is activated by reduction of the plastoquinone pool sensed by the cytochrome *b*_6_*f* complex (Wollman, 2001). By contrast, a recent study of cyanobacteria revealed that neither the phosphorylation and de-phosphorylation reactions nor the cytochrome *b*_6_*f* complex was involved in the regulation of the state transitions (Calzadilla et al., 2019). In cyanobacteria, at least four processes responsible for the state transitions have been proposed: (1) lateral movement of PBSs between PSII and PSI; (2) energetic decoupling of PBSs from either or both of PSII and PSI; (3) connection of PSII and PSI followed by exciton transfer from PSII to PSI (also called energy spillover); (4) intrinsic PSII quenching located in PSII reaction center proteins and/or PSII core antenna proteins (Calzadilla and Kirilovsky, 2020). Recent studies suggested that the cyanobacterial qT was dominantly attributed to intrinsic PSII quenching and partially attributed to energetically decoupled PBSs (Bhatti et al., 2020; Bhatti et al., 2021). Notably, intrinsic PSII quenching was proposed to be involved in both cyanobacterial qT and plant qE, suggesting that different components of NPQ classified by their kinetics and triggering factors may share a fluorescence quenching process in common.

Red algae are a large group of algae displaying unique antenna combination of a eukaryotic type of LHCs specifically bound to PSI and a cyanobacterial type of PBSs (Gardian et al., 2007), and the NPQ mechanisms in red algae appear to be different from those in cyanobacteria, green algae, and plants. None of the proteins in red algae is homologous to the orange carotenoid protein that mediates blue light-driven NPQ in cyanobacteria. A qT-like NPQ modulated by illumination of different colors was identified in the unicellular red alga *Porphyridium cruentum* using 77K fluorescence emission spectroscopy (Murata, 1969). Two mutually non-exclusive processes, energy spillover and energetic decoupling of PBSs from PSII, were proposed to contribute to the qT-like NPQ based on a strong decrease of the fluorescence emission from PSII and a slight increase of the fluorescence emission from PSI (Murata, 1969). The energy spillover model was further supported by an increased photochemical rate of PSI in *P. cruentum* (Ley and Butler, 1977) and reported in the multicellular red algae *Chondrus crispus* (Kowalczyk et al., 2013). Instead of the qT-like NPQ, a qE-like NPQ was proposed in *P. cruentum* and *Rhodella violacea* (Delphin et al., 1996, 1998). Intrinsic PSII quenching was considered as the process that accounted for this qE-like NPQ, since PSII fluorescence decreased while PSI and PBS fluorescences were not altered at 77K (Delphin et al., 1996). In addition to the energy spillover and intrinsic PSII quenching, high mobility of PBSs demonstrated by the kinetics of fluorescence recovery after photobleaching (FRAP) suggested that lateral movement of PBSs was involved in the state transitions in *R. violacea* and *P. cruentum* (Liu et al., 2009; Kaňa et al., 2014). The proposition of various single-process models in *R. violacea* and *P. cruentum* highlights the possibility of more than one fluorescence quenching processes contributing to NPQ. It is interesting to note that the proton ionophore nigericin, which was generally used to determine the presence of qE, also suppressed the energy spillover related to the qT-like NPQ (Kowalczyk et al., 2013). Furthermore, 3-(3,4-dichlorophenyl)-1,1-dimethylurea (DCMU) and 2,5-dibromo-6-isopropyl-3-methyl-1,4-benzoquinone (DBMIB) are common photosynthetic inhibitors to facilitate oxidation and reduction, respectively, of the plastoquinone pool. The opposite effects of DCMU and DBMIB on fluorescence quenching, which was generally taken as evidence for qT, also took place in *R. violacea*, in which qE-like NPQ was proposed (Delphin et al., 1996, 1998). These results indicated that in addition to the transmembrane proton gradient and the redox state of the plastoquinone pool, other factors might also be taken into consideration when assessing the modulation of red algal NPQ.

This work aimed at identifying presumably multiple processes of NPQ in the extremophilic red alga *Cyanidioschyzon merolae*. Intrinsic PSII quenching was proposed to contribute to a qE-like NPQ based on variation of the fluorescence properties under different external pH conditions in isolated PSII complexes (Krupnik et al., 2013). No mobile PBSs were observed in another extremophilic red alga *Cyanidium caldarium* using FRAP analysis, suggesting that the state transitions were not involved in NPQ (Kaňa et al., 2014). However, a slight conformational change of PBSs below the optical resolution (∼200 nm) and corresponding to energetic decoupling of PBSs could not be identified by using FRAP analysis. Energy spillover from PSII to PSI in a dark-acclimated state was identified based on the time-resolved fluorescence spectra of *C. merolae* (Ueno et al., 2017), yet whether the degree of energy spillover was enhanced by illumination was not clear.

We utilized several fluorescence analytical methods with light preferentially absorbed by either PBSs or photosystems to assess different fluorescence quenching processes. PBSs consist of phycobiliproteins that absorb light in the green and amber ranges of the spectrum whereas photosystems contain chlorophylls and carotenoids that dominantly absorb light in the blue and red ranges of the spectrum. Intrinsic PSII quenching and energy spillover were evaluated by measuring illumination-induced changes in the fluorescence emitted from PSII and PSI with excitation of photosystems at 77K. Energetic decoupling of PBSs from PSII was assessed by the difference between the functional antenna size of PSII with excitation of PBSs and that with excitation of PSII as well as by the difference between the NPQ level estimated with excitation of PBSs and that estimated with excitation of PSII. Both assessment methods had been employed to identify the energetic decoupling in cyanobacteria (Koblížek et al., 1998; McConnell et al., 2002). Factors that modulated the identified fluorescence quenching processes were further discussed based on how the NPQ kinetics were affected by protein crosslinkers, osmolytes, ionophores, and photosynthetic inhibitors.

## RESULTS

### NPQ kinetics under various illumination and temperature conditions

Conditions suitable for assessing presumably multiple NPQ processes were determined by fluorescence probed using amber detecting light under continuous red illumination with various photosynthetic photon flux densities (PPFDs) and illumination periods at 40°C. Fluorescence quenched to a light-acclimated level was achieved by illuminating algae for 6 min (Supplementary Fig. S1). Two distinct phases, one rapid increment phase followed by one variable decay phase, were identified during the illumination period (Supplementary Fig. S1A). Under illumination in the range from 40 to 300 μmol photons m^-2^ s^-1^, the NPQ level increased at a similar rate in the initial phase. In contrast, a later onset and smaller extent of the NPQ decay was observed under illumination at a higher PPFD, leading to the increase of the light-acclimated NPQ level along with the increase of PPFD. NPQ decreased and approached the dark-acclimated level in 10 min of darkness. The extents of NPQ increment and decay under low illumination at 10 μmol photons m^-2^ s^-1^ were lower than those under the illumination at 40 μmol photons m^-2^ s^-1^, resulting in relatively high NPQ levels after 6 min of illumination at 10 μmol photons m^-2^ s^-1^ and after illumination followed by 10 min of darkness. An insufficient illumination period of 30 s or 1 min led to a relatively high NPQ level after 6-min illumination and 10-min darkness regardless of PPFDs (Supplementary Fig. S1C-D). Illumination for 6 min at 300 and 10 μmol photons m^-2^ s^-1^ were chosen as conditions for further assessment of multiple NPQ processes, and representative fluorescence traces under illumination at 300 and 10 μmol photons m^-2^ s^-1^ at 40°C were shown in Supplementary Fig. S2.

To facilitate the comparison between the NPQ level and the following analysis results, a suitable temperature allowing for the retardation of dynamic processes of NPQ was selected from 40, 34, and 28°C. NPQ kinetics at 34°C was slower than those at 40°C, and the light-acclimated NPQ levels and the following NPQ level after darkness treatment were similar (Fig. 1). The slowest NPQ response was observed at 28°C, yet NPQ decreased to a relatively high level after 30-min darkness following illumination at 300 μmol photons m^-2^ s^-1^ (Fig. 1A). The temperature condition at 34°C was therefore selected for the following analyses.

**Figure 1.**
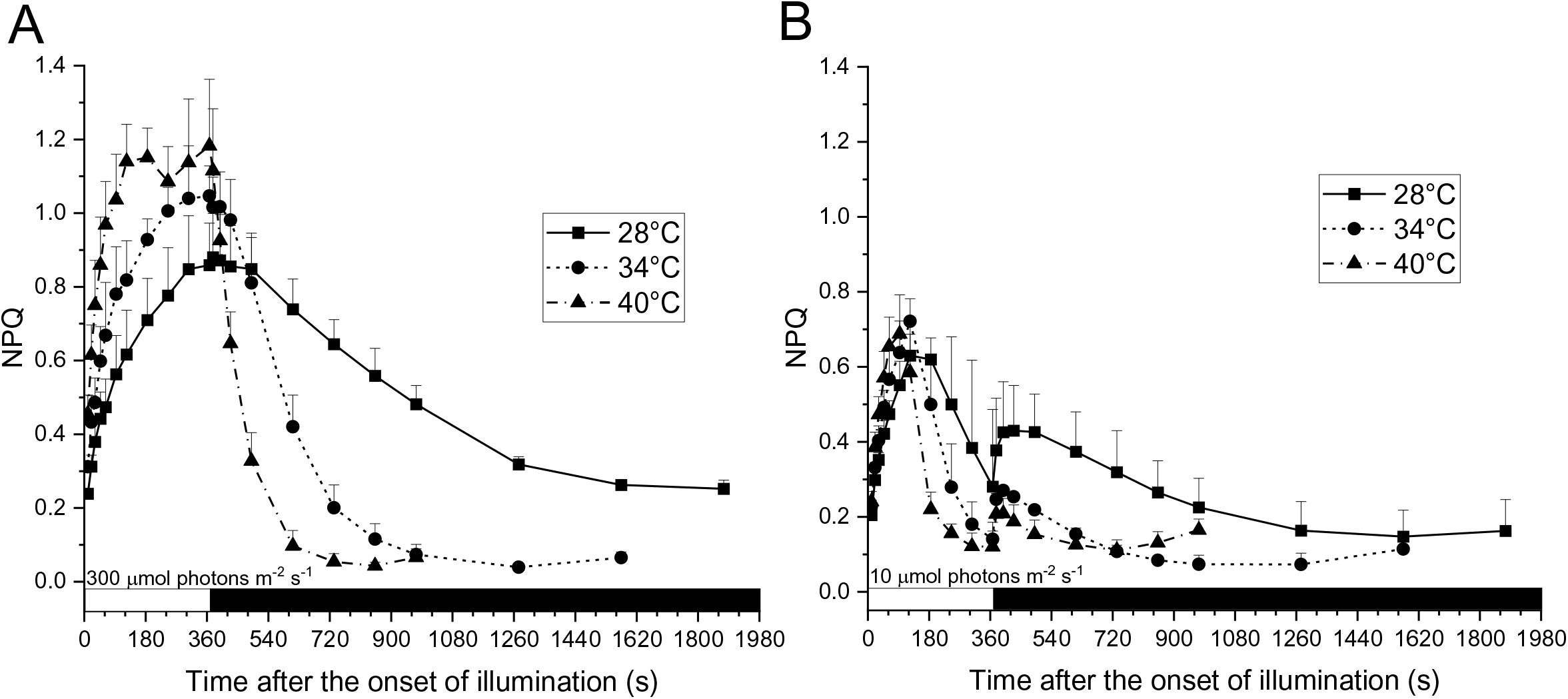
NPQ kinetics at various temperatures. The NPQ level was measured using amber detecting light under continuous red illumination at 300 (A) or 10 μmol photons m^-2^ s^-1^ (B) with illumination periods of approximate 6 min followed by darkness. The white horizontal bar indicates the period of illumination, and the dark horizontal bar indicates the period of darkness. Data are expressed as the average + SD of three independent experiments.

### Analysis of 77K fluorescence emission spectra

To probe transfer of excitons arising from light absorbed by pigments of photosystems or PBSs, 77K fluorescence emission spectra were recorded with the excitation wavelength of 440 nm or 589 nm, respectively. Wavelength-dependent fluorescence amplitudes were normalized by addition of Rhodamine 6G (for 440-nm excitation light) or acid-coated water soluble 900 nm PbS/CdS quantum dots (for 589-nm excitation light; see Materials and Methods for details). The fluorescence spectra were measured after 30 s, 3 min, and 6 min of illumination to assess changes in the exciton transfer in the increment and decay phases of NPQ. Representative Gaussian curves fitted to the fluorescence emission spectrum were shown in Supplementary Fig. S3, and the fluorescence emission components were assigned and listed in Table 1.

**Table 1.** Assignment of fluorescence emission components deconvoluted from the 77K fluorescence emission spectra. Data are expressed as the average ± SD of all the fluorescence spectra (*n* = 54) used in Fig. 2. FWHM, full width at half maximum.

| Name | Assignment | 440-nm excitation |  | 589-nm excitation |  |
| --- | --- | --- | --- | --- | --- |
|  |  | Wavelength (nm) | FWHM (nm) | Wavelength (nm) | FWHM (nm) |
| F660 | PBS | - | - | $659.5 \pm 0.3$ | $28.3 \pm 0.3$ |
| F685 | CP43 | $685.5 \pm 0.4$ | $14.2 \pm 1.2$ | | |
| | CP43 and PBS | | | $685.5 \pm 0.1$ | $13.3 \pm 0.6$ |
| F695 | CP47 | $695.6 \pm 0.3$ | $6.2 \pm 0.3$ | $696.5 \pm 0.2$ | $8.5 \pm 0.3$ |
| F728 | PSI | $728.5 \pm 0.2$ | $29.5 \pm 0.8$ | $728.0 \pm 0.2$ | $31.3 \pm 0.8$ |

Excitation light at 440 nm gave rise to contrasting low fluorescence emitted from CP43 and CP47 of PSII (corresponding to F685 and F695, respectively) and high fluorescence emitted from PSI (corresponding to F728), the latter of which was expected based on a large portion of chlorophyll and carotenoid molecules located in LHCs and energetically coupled to PSI (Fig. 2A). Illumination at 300 and 10 μmol photons m^-2^ s^-1^ led to an overall increase in F728 and decrease in F685 and F695 (Fig. 2A-B). The extent of decrease in F685 and F695 was high and statistically significant, whereas the extent of increase in F728 was low and insignificant (Fig. 2E). Such distinct extents of changes in fluorescence emitted from PSII and PSI suggested that intrinsic PSII quenching accounted for a major portion of NPQ with 440-nm excitation light while energy spillover from PSII-to-PSI played a minor role. It was noted that fluorescence emitted from PSII apparently increased when algae were illuminated from 3 to 6 min at 10 μmol photons m^-2^ s^-1^ (Fig. 2B and 2E), consistent with the decreases in the NPQ level in the decay phase (Fig. 1B). In the darkness for 25 min after illumination, F685, F695, and F728 approached their dark-acclimated levels, corresponding to the low NPQ level under the same experimental condition (Fig. 1, 2A-B, and 2E).

**Figure 2.**
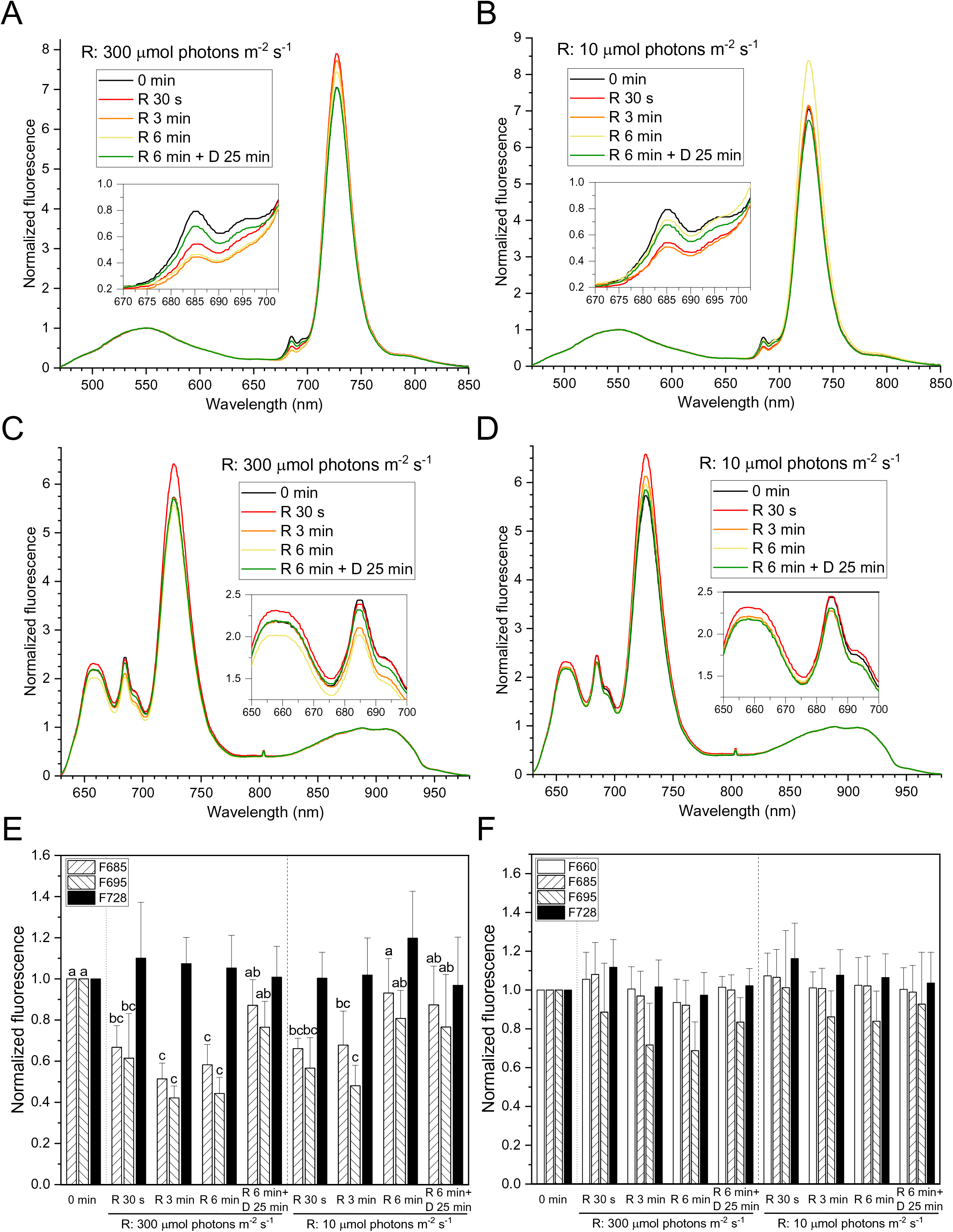
Changes in the 77K fluorescence emission spectrum under illumination. (A)-(D) Changes in the fluorescence emission spectrum with the excitation wavelength of 440 nm (A, B) or 589 nm (C, D). The fluorescence emission spectrum was measured after continuous red illumination at 300 (A, C) or 10 μmol photons m^-2^ s^-1^ (B, D) for 30 s, 3 min, 6 min, or after 25 min of darkness following the 6-min illumination. The fluorescence emission spectrum was normalized to the maximum fluorescence level of the exogenous fluorophore [Rhodamine 6G for (A), (B) and PbS/CdS quantum dots for (B), (D)] as 1. Data are expressed as the average of six independent experiments. (E)-(F) Kinetics of the amplitudes of fluorescence emission components with the excitation wavelength of 440 nm (E) or 589 nm (F). The amplitude of each fluorescence emission component was normalized and compared to that in the dark-acclimated state (i.e. 0 min). Data are expressed as the average + SD of six independent experiments. Lowercase letters indicate significantly different groups as determined using one-way ANOVA followed by Tukey’s multiple comparison test. R, red illumination; D, darkness.

Light at 589 nm was predominantly absorbed by phycobilins and gave rise to fluorescence emitted from PBS (corresponding to F660) in addition to F685, F695, and F728 (Fig. 2C-D). A high peak amplitude ratio of F685 over F695 with 589-nm excitation light compared to the ratio with 440-nm excitation light was attributed to both the PBS terminal emitter and PSII-CP47 contributing to F685 (Scott et al., 2006). Illumination induced decreases in F695 with the 589-nm excitation light like those with the 440-nm excitation light (Fig. 2E-F). The decrease in F695 with excitation of PBSs was expected to be equivalent or enhanced in the absence or presence, respectively, of energetic decoupling of PBSs from PSII. However, the extent of decreases in F695 with 589-nm excitation light was lower than that with 440-nm excitation light (Fig. 2E-F). The dampening effect of the changes in the PSII fluorescence with 589-nm excitation light might be due to the impact of freezing on the connection between the PBS and PSII (see Discussion about interpretation of the changes in F660 and F695). Therefore, whether energetic decoupling of PBSs occurred under illumination could not be confirmed by the fluorescence emission spectra with 589-nm excitation light.

### Changes in the functional antenna sizes of PSII with excitation of PBS and PSII

As an alternative method to assess the involvement of energetic decoupling of PBSs from PSII in NPQ, the functional antenna size of PSII with excitation of PBSs using amber actinic light and that with excitation of PSII using blue actinic light were compared. The functional antenna size of PSII was calculated as the reciprocal of the proportion of the area above the fluorescence rise curve in the presence of DCMU (see Materials and Methods for details). The functional antenna size decreased due to either a reduced number of antenna pigments energetic coupled to the PSII reaction center or enhanced quenching of excitons, as competition of quenchers against exciton trapping for the reduction of Q_A_ was proposed as a cause of the lowering of the fluorescence level during the fluorescence rise in the presence of DCMU (Belgio et al., 2014; Tian et al., 2019). When exitonic flux within PSII was deviated to a quencher within PSII (i.e. intrinsic PSII quenching) or to the PSI reaction center (i.e. energy spillover), the functional antenna size with excitation of PBSs proportionally decreased with respect to the functional antenna size with excitation of PSII. When photons absorbed by PBSs for PSII photochemistry and/or excitonic transfer efficiency from PBSs to PSII decreased (i.e. energetic decoupling of PBSs from PSII), the functional antenna size with excitation of PBSs was expected to decrease while the functional antenna size with excitation of PSII was expected to be unchanged. The degree of energetic decoupling could therefore be assessed by employing the change in the difference between the two functional antenna sizes as an estimate, especially when the two functional antenna sizes were known to be additionally reduced by the intrinsic PSII quenching and/or the energy spillover.

As shown in Fig. 3, the decrease in the functional antenna sizes of PSII with excitation of PBSs and PSII already took place upon illumination for 10 s. Reduction in the antenna size with excitation of PSII was associated with the intrinsic PSII quenching accounting for the major portion of NPQ, as determined by the 77K fluorescence spectra (Fig. 2A and E). The extent of decreases in the antenna size with excitation of PBSs was much higher than that with excitation of PSII, indicating that PBSs were energetically decoupled from PSII. The difference between the two functional antenna sizes was persistently smaller at 300 μmol photons m^-2^ s^-1^ than that at 10 μmol photons m^-2^ s^-1^ (Fig. 3C-D), suggesting that the degree of energetic decoupling increased over PPFD. The low functional antenna sizes with excitation of PBSs or PSII corresponded to the high NPQ levels under illumination except in the NPQ decay phase under illumination at 10 μmol photons m^-2^ s^-1^ (Fig. 1B and 3D). The inconsistency between the changes in the functional antenna size and the changes in the NPQ level could be attributed to additional processes that steadily enhanced fluorescence regardless of the redox state of Q_A_.

**Figure 3.**
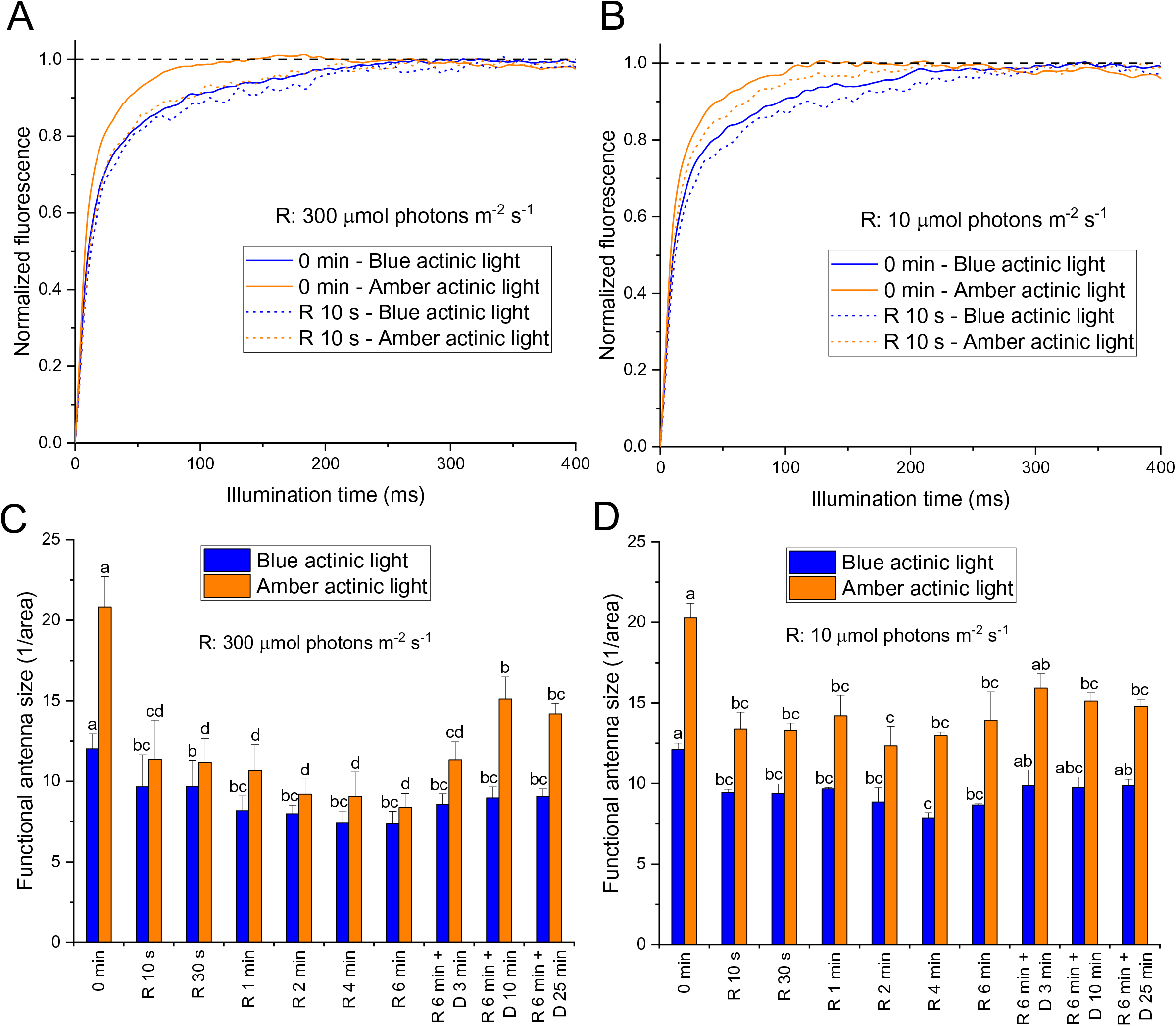
Changes in the functional antenna size of PSII under illumination. (A)-(B) Averaged fluorescence induction curve in the dark-acclimated state (0 min) and after 10 s of red illumination at 300 (A) or 10 μmol photons m^-2^ s^-1^ (B). The fluorescence induction curve was measured 20 s after the addition of DCMU under illumination of blue or amber actinic light at 200 μmol photons m^−2^ s^−1^. The values of fluorescence induction curve were normalized to 0 equivalent to the fluorescence level before actinic light and to 1 equivalent to the maximum fluorescence level induced actinic light. The curves are expressed as the average of six independent experiments. The averaged fluorescence induction curves after the other time courses of illumination and darkness treatment are shown in Supplementary Fig. S4-S5. (C)-(D) Functional antenna size of PSII estimated from the fluorescence induction curve in the presence of DCMU. The fluorescence induction curve was measured after different time courses of illumination and darkness treatment. The functional antenna size was calculated as the reciprocal of the proportion of the area above the fluorescence rise curve. Data are expressed as the average + SD of six independent experiments. Lowercase letters indicate significantly different groups as determined using one-way ANOVA followed by Tukey’s multiple comparison test. R, red illumination; D, darkness.

### NPQ kinetics measured with detecting lights of different colors

To assess the contribution of energetic decoupling of PBSs from PSII to NPQ, fluorescence was probed by using green, blue, and amber detecting lights sequentially and repeatedly (see Materials and Methods for details). Energetic decoupling of PBSs from PSII was confirmed by the result that the NPQ level measured using amber detecting light (preferentially exciting PBSs) was larger than that measured using green detecting light (exciting PBSs in a large part and PSII in a small part) or blue detecting light (preferentially exciting PSII; Fig. 4A-B). The onset of energetic decoupling was as rapid as other fluorescence quenching processes, since the NPQ level measured using amber or green detecting light was already higher than that measured using blue detecting light at 10 s of illumination. A linear curve was fit to the NPQ level measured using either amber or green detecting light versus the NPQ level measured using blue detecting light throughout the experiment, and the linear correlation ratio at 300 μmol photons m^-2^ s^-1^ was higher than that at 10 μmol photons m^-2^ s^-1^ (Supplementary Fig. S6). Such a light-dependent ratio in the NPQ levels was in line with light-dependent difference in the functional antenna sizes (Fig. 3C-D) and thus considered as another estimate for assessing the degree of energetic decoupling.

**Figure 4.**
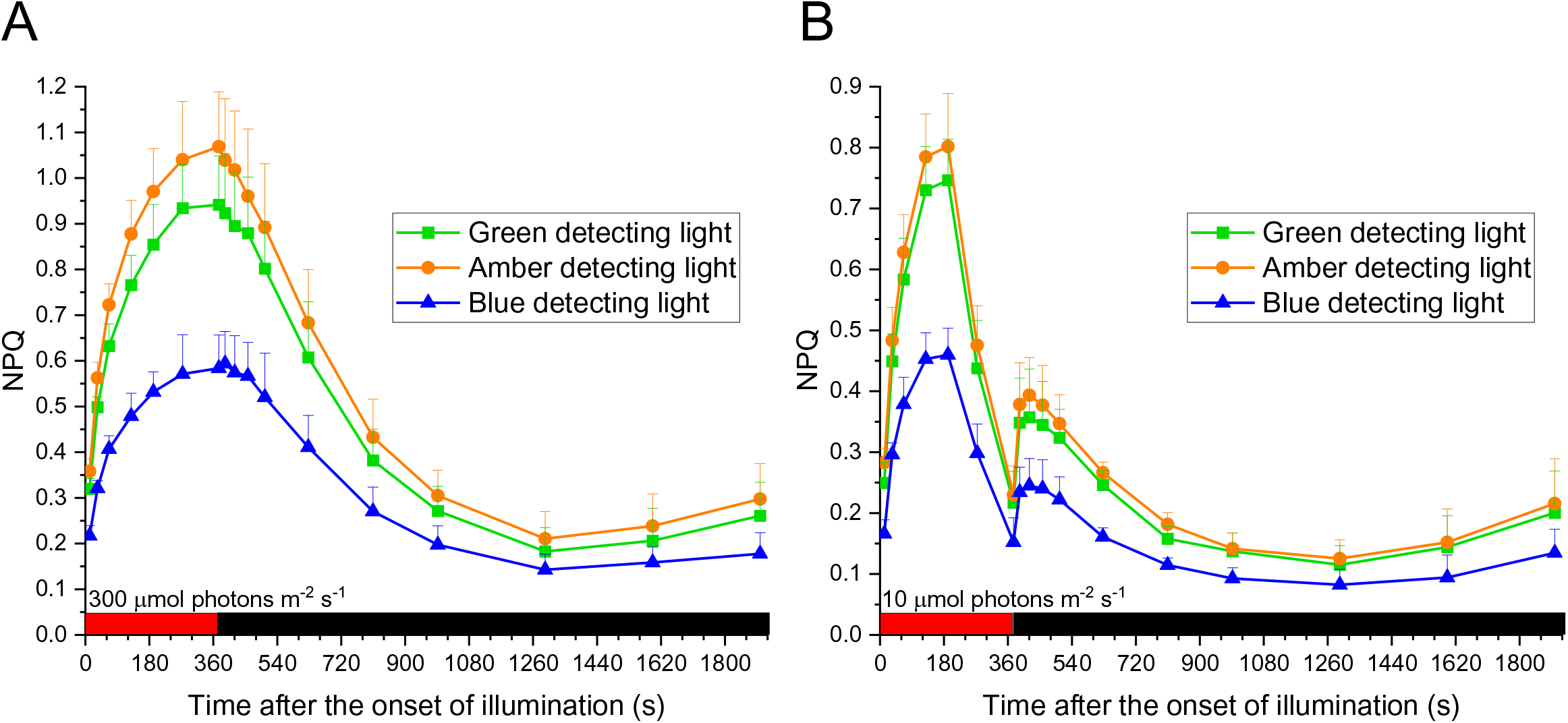
NPQ kinetics using detecting lights of different colors under illumination. The NPQ level was estimated from fluorescence probed using green, blue, and amber detecting lights sequentially and repeatedly. Data are expressed as the average + SD of three independent experiments. The red horizontal bar indicates the period of illumination, and the dark horizontal bar indicates the period of darkness.

It was previously reported that a saturating multi-turnover light pulse was sufficient to trigger fluorescence quenching in red algae (Delphin et al., 1998; Kowalczyk et al., 2013). As the energetic decoupling of PBSs from PSII and the other fluorescence quenching processes were simultaneously induced by 10 s of continuous illumination, the energetic decoupling was expected to be driven by light pulses. Further verification was performed by measuring the NPQ kinetics in the presence of multi-turnover light pulses applied with a 2-s, 10-s, or 30-s interval of darkness. The extent of increase in the NPQ level after each pulse was enhanced over the dark interval (Supplementary Fig. S7A). The energetic decoupling of PBSs from PSII, as determined by reduction in the difference between the functional antenna size of PSII with excitation of PBSs and that with excitation of PSII, was already observed after one light pulse (Supplementary Fig. S7B). The degree of energetic decoupling was similar with that under illumination at 10 μmol photons m^-2^ s^-1^ (Fig. 3D and Supplementary Fig. S7B). By contrast, the degree of intrinsic PSII quenching, as determined by the extent of decrease in the functional antenna size with excitation of PSII, was less significant than that under illumination (Fig. 3C-D, and Supplementary Fig. S7B). Irrespective of the interval of darkness, the degrees of energetic decoupling remained similar after its onset, as estimated from the difference between the two functional antenna sizes of PSII and from the ratio of the NPQ level measured using amber detecting light over that measured using blue detecting light (Supplementary Fig. S7B-D). The similar degrees of energetic decoupling in multiple light pulses conditions were consistent with those in continuous illumination conditions. On the other hand, the NPQ level related to the other fluorescence quenching processes was dynamic depending on the illumination period and intensity.

### Effect of glutaraldehyde, betaine, and glycerol on the energetic decoupling of PBSs from PSII

To evaluate whether the energetic decoupling of PBSs from PSII could be mechanistically distinguished from the other fluorescence quenching processes, the effect of two types of chemicals on the NPQ kinetics was analyzed. Protein crosslinkers, such as glutaraldehyde, were reported to stabilize protein structure by crosslinking two reactive groups of amino acids (Scott et al., 2006). Stabilizing osmolytes, such as betaine and glycerol, were proposed to induce protein folding to a stable conformation (Rösgen et al., 2005). Different reactions of these two types of chemicals were illustrated by 77K fluorescence emission spectra. Whereas two osmolytes, betaine and glycerol, suppressed the fluorescence emitted from PBS, PSII, and PSI with 440-nm and 589-nm excitation lights in a concentration-dependent manner, the crosslinker glutaraldehyde did not significantly affect the fluorescence amplitude (Supplementary Fig. S8). It was noted that betaine and glycerol suppressed both F685 and F695 with excitation of photosystems yet suppressed only F685 with excitation of PBSs (Supplementary Fig. S8F). The resulting enhancement of F695 with excitation of PBSs suggested that connection between the PBS and PSII was disturbed by betaine or glycerol. On the other hand, no prominent effect of glutaraldehyde on the connection between the PBS and PSII was observed. Regardless of the different reactions shown by the 77K fluorescence emission spectra, inhibition of light-triggered NPQ was observed when either glutaraldehyde or betaine was added in the dark-acclimated state (Fig. 5). It was noted that 0.025% of glutaraldehyde or 283 mM of betaine was sufficient to completely prevent the onset of NPQ and thus considered as the effective concentration. Distinct effects on NPQ kinetics were observed when glutaraldehyde or betaine was added immediately after 3 min of illumination (Fig. 6). While sub-effective concentration (0.005%) of glutaraldehyde lowered the following NPQ levels, 0.025% of glutaraldehyde kept NPQ to the level right before the addition of glutaraldehyde throughout the experiment (Fig. 6A-B). In contrast to glutaraldehyde, 56.6 mM of betaine did not effectively alter the NPQ kinetics, and 283 mM of betaine prevented the NPQ level from decay (Fig. 6C-D). Strikingly, the degree of energetic decoupling, as determined by the difference between the NPQ level measured using amber or green detecting light and that measured using blue detecting light, was elevated by betaine (Fig. 6D). The effect of glycerol on the NPQ kinetics was comparable with that of betaine (Supplementary Fig. S9), indicating that the energetic decoupling of PBSs from PSII was specifically affected by stabilizing osmolytes that disturbed the conformation associated with the connection between the PBS and PSII.

**Figure 5.**
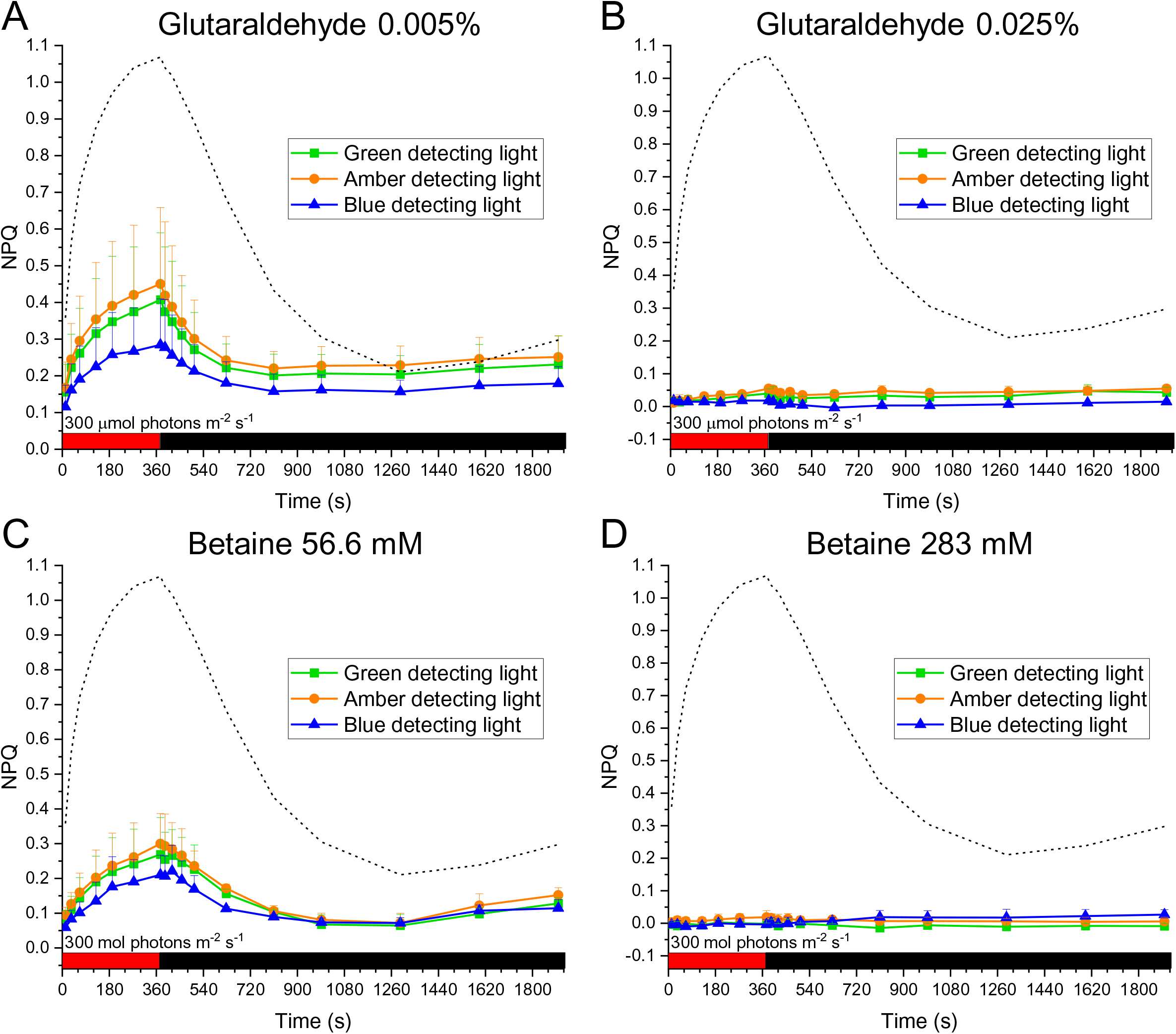
Effect of glutaraldehyde or betaine added in the dark-acclimated state on the NPQ kinetics using detecting lights of different colors. (A)-(B) The NPQ kinetics in the presence of 0.0005% (A) and 0.025% (B) of glutaraldehyde added 20 min before experiments under illumination at 300 μmol photons m^−2^ s^−1^. (C)-(D) The NPQ kinetics in the presence of 56.6 mM (C) and 283 mM (D) of betaine added 20 min before experiments under illumination at 300 μmol photons m^−2^ s^−1^. The NPQ level was measured using green, blue, and amber detecting lights sequentially and repeatedly. Data are expressed as the average + SD of three independent experiments. The dash line indicates the NPQ kinetics of control in the absence of glutaraldehyde or betaine using amber detecting light. The red horizontal bar indicates the period of illumination, and the dark horizontal bar indicates the period of darkness.

**Figure 6.**
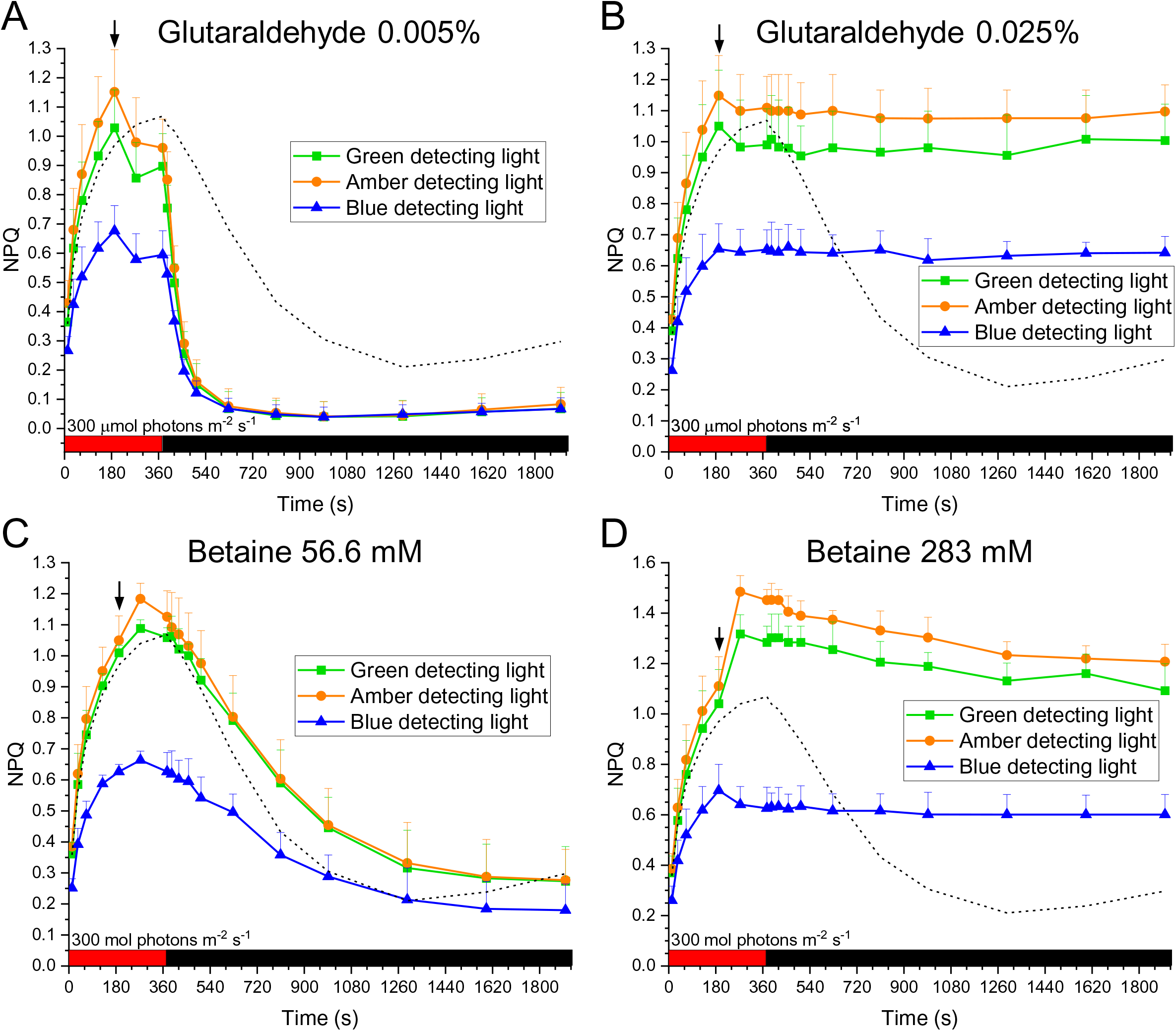
Effect of glutaraldehyde or betaine added in the light-acclimated state on the NPQ kinetics using detecting light of different colors. (A)-(B) The NPQ kinetics in the presence of 0.0005% (A) and 0.025% (B) of glutaraldehyde added at 3 min of illumination at 300 μmol photons m^−2^ s^−1^. (C)-(D) The NPQ kinetics in the presence of 56.6 mM (C) and 283 mM (D) of betaine at 3 min of illumination at 300 μmol photons m^−2^ s^−1^. The NPQ level was measured using green, blue, and amber detecting lights sequentially and repeatedly. Data are expressed as the average + SD of three independent experiments. The dash line indicates the NPQ kinetics of control in the absence of glutaraldehyde or betaine using amber detecting light. The arrow marks the time at which glutaraldehyde or betaine is added. The red horizontal bar indicates the period of illumination, and the dark horizontal bar indicates the period of darkness.

### Effect of ionophores on the NPQ kinetics measured with detecting lights of different colors

The transmembrane proton gradient was reported to account for NPQ in *C. merolae* (Krupnik et al., 2013). In addition, electrostatics of photosystems was proposed to modulate red algal NPQ (Kowalczyk et al., 2013). As electrostatic field distribution around the thylakoid membrane was influenced by transmembrane electrochemical gradient of ions, further assessment and verification were conducted by addition of various kinds of ionophores including nigericin (dissipating concentration gradient of protons by exchanging K^+^ or Na^+^ for H^+^ across the membrane), nonactin (dissipating electrochemical gradient of some cations across the membrane), carbonyl cyanide m-chlorophenyl hydrazone (CCCP) and triclosan (both dissipating electrochemical proton gradient across the membrane). All these ionophores affected the NPQ kinetics to different extents (Fig. 7A-B). Under illumination at 300 μmol photons m^-2^ s^-1^, NPQ was largely suppressed by nigericin and became completely diminished by triclosan (Fig. 7A), suggesting that the transmembrane gradient of protons played a major role in the mediation of induction of NPQ while the transmembrane gradient of other ions played a minor role. The role of transmembrane gradient of ions other than protons in modulating NPQ was also supported by the enhancement effect of nonactin on the NPQ level in the absence and the presence of nigericin (Fig. 7A). The contribution of the ions other than protons to NPQ could be neglected under illumination at 10 μmol photons m^-2^ s^-1^, as both nigericin and triclosan completely diminished NPQ (Fig. 7B). Nevertheless, nonactin lowered the NPQ level and abolished the NPQ decay phase under illumination at 10 μmol photons m^-2^ s^-1^ (Fig. 7B). The contrasting enhancement and suppression effects of nonactin on NPQ kinetics could be due to different electrochemical gradients of cations across the membrane under illumination at 300 and 10 μmol photons m^-2^ s^-1^, respectively (see Discussion). Notably, nigericin and/or nonactin facilitated the relaxation of NPQ to the dark-acclimated level during the darkness treatment after illumination regardless of their different effects on the NPQ kinetics under illumination (Fig. 7A-B).

**Figure 7.**
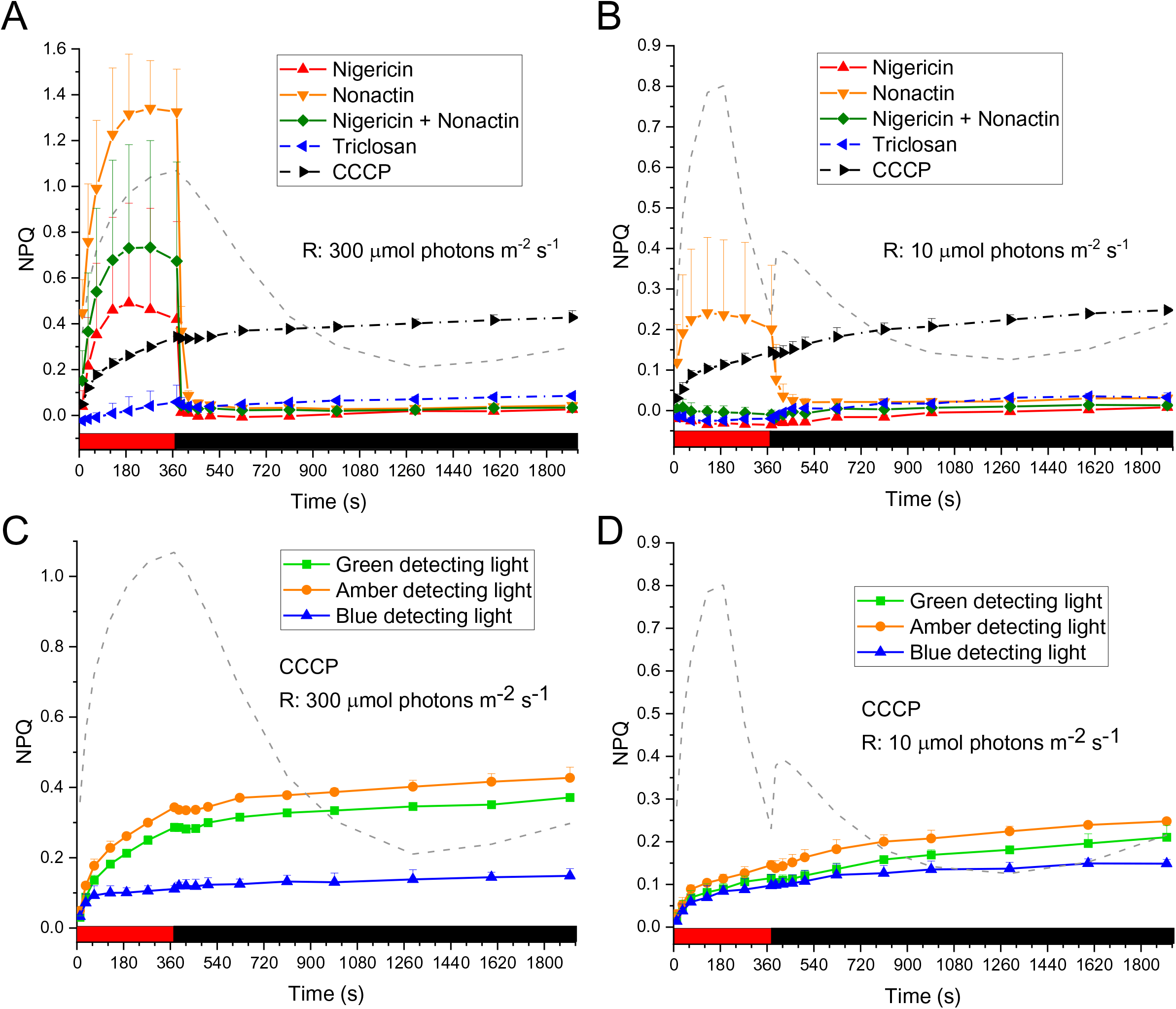
Effect of ionophores on the NPQ kinetics. (A)-(B) The NPQ kinetics in the presence of ionophores under illumination at 300 (A) and 10 (B) μmol photons m^−2^ s^−1^. The ionophores were added 20 min before experiments, and the NPQ level was measured using amber detecting light. (C)-(D) The NPQ kinetics in the presence of CCCP under illumination at 300 (A) and 10 (B) μmol photons m^−2^ s^−1^. The ionophores were added 20 min before experiments, and the NPQ level was measured using green, blue, and amber detecting lights sequentially and repeatedly. Data are expressed as the average + SD of three independent experiments. The dash line indicates the NPQ kinetics of control in the absence of ionophores using amber detecting light. The red horizontal bar indicates the period of illumination, and the dark horizontal bar indicates the period of darkness.

Unexpectedly, CCCP slowly enhanced NPQ and displayed an effect distinct from triclosan (Fig. 7A-B), although both chemicals were reported to dissipate electrochemical proton gradient across the membrane. Strikingly, CCCP enhanced the ratio between the NPQ level measured with amber detecting light and that measured with blue detecting light under illumination at 300 μmol photons m^-2^ s^-1^ (Supplementary Fig. S10A). Accordingly, the enhancement of the NPQ level measured using amber or green detecting light was more prominent at 300 μmol photons m^-2^ s^-1^ than that at 10 μmol photons m^-2^ s^-1^, whereas the NPQ level measured using blue detecting light was steadily low (∼0.1) under both illumination conditions (Fig. 7C-D). The differential enhancement of the NPQ levels measured with detecting lights of different colors was indicative of the energetic decoupling selectively affected by CCCP. In search of how CCCP influenced the energetic decoupling, it was noted that the operating efficiency of PSII photochemistry (Φ_PSII_) was particularly decreased in the presence of CCCP, as compared to other ionophores, during the period of darkness after illumination in a light-dependent manner (Supplementary Fig. S10C-D). The effect of CCCP on Φ_PSII_ was consistent with the inhibition of oxygen evolution that was caused by oxidation of CCCP at the PSII donor side and intensified under illumination (Homann, 1971). Addition of DCMU that blocked the electron flow to Q_B_ and the plastoquinone pool counteracted the effect of CCCP on the degree of energetic decoupling (Fig. 8A). Combination of triclosan and DBMIB led to progressive increases in the NPQ level yet did not change the degree of energetic decoupling, the result of which was similar to that in the presence of CCCP + DCMU (Fig. 8A-B). Notably, DCMU and DBMIB had opposite effects on the redox state of plastoquinone pool, yet DBMIB did not enhance the degree of energetic decoupling. Therefore, the counteracting effect of DCMU on CCCP-induced energetic decoupling was unlikely mediated by the redox state of the plastoquinone pool. Furthermore, the NPQ levels measured with green, blue, and amber detecting lights was proportionally suppressed by either DCMU or DBMIB alone to a slightly different extent (Fig. 8C-D). The overall results with addition of DCMU or DBMIB suggested that the redox state of the plastoquinone pool did not play a significant role in the regulation of multiple NPQ processes including the intrinsic PSII quenching and the energetic decoupling.

**Figure 8.**
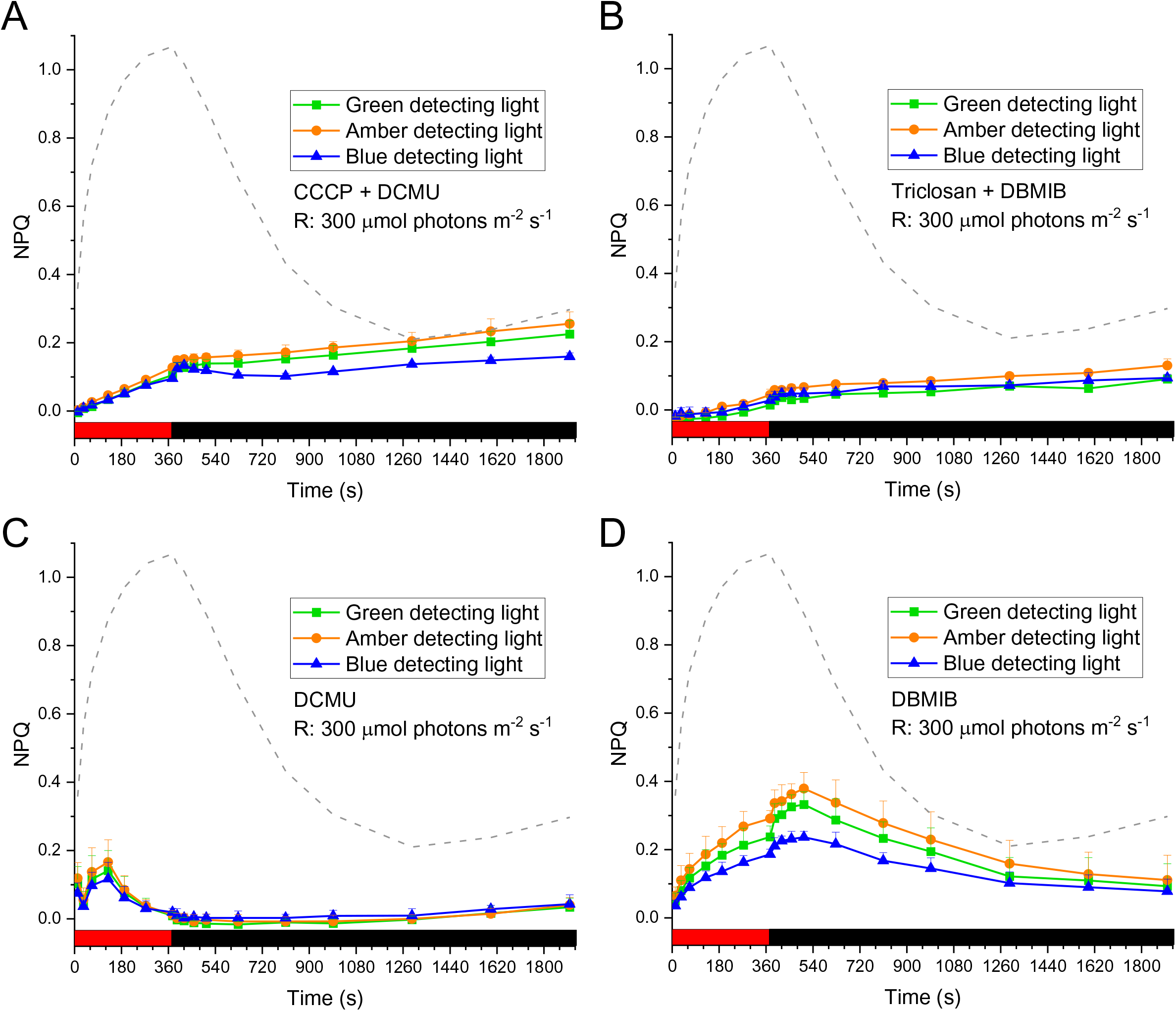
Combination effect of ionophores and photosynthetic inhibitors on the NPQ kinetics using detecting lights of different colors. The NPQ kinetics was measured in the presence of CCCP + DCMU (A), triclosan + DBMIB (B), DCMU (C), and DBMIB (D) under illumination at 300 μmol photons m^−2^ s^−1^. The ionophores were added 20 min before experiments, and the NPQ level was measured using green, blue, and amber detecting lights sequentially and repeatedly. Data are expressed as the average + SD of three independent experiments. The dash line indicates the NPQ kinetics measured using amber detecting in the absence of ionophores and photosynthetic inhibitors. The red horizontal bar indicates the period of illumination, and the dark horizontal bar indicates the period of darkness. R, red illumination.

## DISCUSSION

The present work identified energetic decoupling of PBSs from PSII involved in NPQ with excitation of PBSs in the extremophilic red alga *C. merolae*. Furthermore, we confirmed that intrinsic PSII quenching contributed to NPQ as previously reported (Krupnik et al., 2013). The two fluorescence quenching processes could be distinguished by their kinetics and reactions with protein crosslinkers, osmolytes, ionophores, and photosynthetic inhibitors. Both the energetic decoupling and the intrinsic PSII quenching reacted rapidly in response to light. Whereas the energetic decoupling remained steady after its induction, the intrinsic PSII quenching was dynamic depending on the illumination period and intensity. Stabilizing osmolytes added under illumination enhanced the degree of energetic decoupling yet did not affect the degree of intrinsic PSII quenching. Furthermore, the energetic decoupling was specifically induced by CCCP under high illumination. Our results provided valuable information for discussion of mechanisms accounting for the modulation of NPQ in red algae.

The energetic decoupling of PBSs from PSII was successfully identified by estimation of the functional antenna size and the NPQ level measured with light preferentially absorbed by PBSs or PSII, yet it could not be clearly observed by fluorescence emission spectroscopy with excitation of PBSs at 77K. A slight increase in the PBS fluorescence (i.e. F660) at 30 s of illumination was indicative of energetic decoupling of PBSs, yet the following decreases of the PBS fluorescence under illumination for 3 and 6 min appeared to contradict the energetic decoupling model (Fig. 2F). It was noted that the small extent of decreases in the PSII fluorescence (i.e. F695) with excitation of PBSs did not correspond to a relatively high NPQ level, whereas the change in F695 with excitation of photosystems and the change in the NPQ level were comparable (Fig. 1 and 2). The inconsistent changes in the PSII fluorescence with excitation of PBSs at 34°C and that at 77K might indicate the effect of temperature on the exitonic transfer from PBSs to PSII. Cryogenic temperature was reported to be causing changes in the membrane protein structure (Mehra et al., 2020). As discussed below, the light-induced energetic decoupling likely involves a slight increase in the distance between the PBS and PSII. Such a subtle change in the connection between the PBS and PSII may be more readily affected by freezing than the conformational change around the quenching site associated with the intrinsic PSII quenching.

Two possible sites involved in the energetic decoupling of PBSs from PSII are (1) the chromophores inside PBSs and (2) the interface between the PBS and PSII. The PBS generally comprises three types of phycobiliproteins: phycoerythrin (PE), phycocyanin (PC), and allophycocyanin, all of which are assembled with the assistance of linker polypeptides. Detachment of some PEs from other parts of the PBS was considered as one mechanism underlying the energetic decoupling based on a decrease in the fluorescence emitted from PBSs and a short-term rise of the fluorescence emitted from PE (Liu et al., 2008). *C. merolae* has no PE, and no additional fluorescence emitted from PC was observed in the 77K fluorescence emission spectra (Fig. 2C-D). Therefore, it is more likely that the detachment of PBSs from PSII accounts for the energetic decoupling of PBSs from PSII.

A nearly fixed ratio between the NPQ level measured with amber detecting light and that measured with blue detecting light provides understanding of the energetic decoupling mechanism in accordance with the detachment of PBSs from PSII. The Stern-Volmer relationship is implicitly shown by calculation of the NPQ level in the following formula: NPQ = C_SV_ × [Q], where C_SV_ is the Stern-Volmer quenching constant, and [Q] is the quencher concentration. The Stern-Volmer formula for the NPQ level measured using amber and blue detecting light, can be expressed as NPQ_amber_ = C_SV,amber_ × [Q]_amber_ and NPQ_blue_ = C_SV,blue_ × [Q]_blue_, respectively. The ratio, *k*, between the NPQ level measured with amber detecting light and that measured with blue detecting light is calculated as:

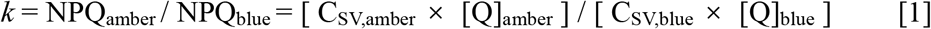

For simplicity, the same quencher concentration corresponding to [Q]_amber_ and [Q]_blue_ is assumed, and thus [Q]_amber_ = [Q]_blue_ = [Q]. By reorganizing [1] we obtain:

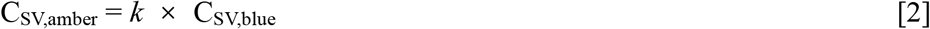

The *k* value is interpreted as an additional loss in the energy transfer efficiency with excitation using amber light compared to that with excitation using blue light and is equivalent to a change in the energy transfer efficiency from PBSs to PSII. The energy transfer efficiency is implicitly explained by Förster resonance energy transfer mechanism and determined by the distance between the PBS and PSII (Krasilnikov et al., 2020). A constant *k* level may therefore reflect a constant distance on average at which PBSs are detached from PSII. If the energy transfer efficiency follows the Förster resonance energy transfer mechanism in which the efficiency is inversely proportional to the sixth power of the distance between the donor and acceptor pigment, an increase of distance by ∼10% is needed for the value of *k* to be 1.8. A slight shift of less than 0.5 nm in distance suffices to induce the energetic decoupling of PBSs from PSII as the shortest distance of the two pigments between the PBS and PSII is ∼4 nm (Chang et al., 2015; Krasilnikov et al., 2020). Such a mechanistic view is compatible with the previous finding that no mobile PBSs between PSII and PSI was observed in another extremophilic red alga *C. caldarium* using FRAP analysis (Kaňa et al., 2014), since PBS movement of less than one nanometer can result in strong energetic decoupling and is not detectable under light microscope.

In contrast to the apparent roles of PBSs and PSII played in red algal NPQ, the involvement of PSI in NPQ is less apparent. The results of 77K fluorescence spectroscopy with excitation at 440 nm compared with those with excitation at 589 nm suggested that a significant portion of light energy absorbed by PBSs was transferred to PSI (Fig. 2). Both direct energy transfer from PBSs to PSI and indirect energy transfer from PBSs to PSI via PSII were proposed using time-resolved fluorescence analysis (Ueno et al., 2017). The direct energy transfer was supported by a partial PBS identified in an isolated PSI complex (Busch et al., 2010), while no functional association of the PBS with PSI was identified using another PSI isolation procedure (Haniewicz et al., 2018). Regardless of direct or indirect energy transfer from PBSs to PSI, whether excitonic transfer toward PSI changes upon induction of NPQ has not been clearly demonstrated. Excitonic connection or disconnection of PBSs to PSI was not clearly demonstrated by the fluorescence emission spectra with excitation of PBSs, yet it may be due to the freezing effect as discussed before. The putative PSII-to-PSI energy spillover induced by illumination was identified based on increase in the PSI fluorescence and decrease in the PSII fluorescence with excitation of photosystems at 77K, yet its contribution to fluorescence quenching was much less than the contribution of intrinsic PSII quenching (Fig. 2A-B). This result was consistent with the finding that energy redistribution between PSII and PSI was neglectable under illumination in the red alga *R. violacea* (Delphin et al., 1996).

The contrasting effects of glutaraldehyde, betaine, and glycerol added between darkness and illumination provide a novel aspect on the fluorescence quenching processes. Glutaraldehyde was proposed to crosslink PBSs with photosystems and thus to deplete PBS mobility (Kaňa et al., 2014). Betaine was proposed to fix PBSs on the thylakoid membrane (Li et al., 2001) or to affect connection of PBSs with PSII and PSI (Papageorgiou et al., 1999; Calzadilla et al., 2019), the latter observation being supported by the analysis result of the 77K fluorescence spectra (Supplementary Fig. S8). All these proposed mechanisms are collectively concerned with protein stabilization of PBSs. Addition of glutaraldehyde, betaine, or glycerol in the darkness suppressed NPQ with excitation of both PBSs and PSII, suggesting that in addition to PBSs, photosystems were also stabilized (Fig. 5). The effect of glutaraldehyde, betaine, or glycerol added under illumination on NPQ appear to contradict the suppression of NPQ caused by the same chemical compound added in the darkness. Since the reactions of different chemical compounds with the photosynthetic apparatus are expected to be the same, the distinct effects of these chemical compounds on the NPQ kinetics imply that protein conformations of the photosynthetic apparatus relevant to NPQ under dark conditions are different from those under illuminated conditions. It is therefore conceivable that the PBS and PSII undergo protein conformational changes corresponding to changes in the NPQ level. With the effective concentration, glutaraldehyde or stabilizing osmolytes (betaine and glycerol) stabilizes the conformations of PBSs and photosystems, thus keeping NPQ at a nearly constant level in the light-acclimated state. The contrasting effects of glutaraldehyde added at the effective concentration between darkness and illumination on the NPQ kinetics were previously observed and proposed to be attributed to the changes in the conformation of photosynthetic apparatus associated with the sites accessible to glutaraldehyde (Scott et al., 2006). The contrasting effect of low and high concentration of glutaraldehyde on depletion and retention of NPQ, respectively, can also be interpreted by the accessibility of glutaraldehyde to proteins. Formation of crosslinking requires an adequate distance between two reactive groups and depends on the concentration of crosslinkers (Barbosa et al., 2014). Based on the NPQ kinetics, glutaraldehyde molecules at low concentration (0.005%) may react with sites that facilitate the transformation of the protein conformations to the dark-acclimated state. At higher concentration (0.025%), a larger number of glutaraldehyde molecules may crosslink more reactive groups of amino acids and thus keep the protein conformations in the light-acclimated state. In all the cases, glutaraldehyde was not able to affect the energetic decoupling of PBSs from PSII, which is consistent with no significant effect on the 77K fluorescence related to the connection between the PBS and PSII (Supplementary Fig. S8). By contrast, the effect of stabilizing osmolytes on the connection between the PBS and PSII was revealed by the changes in the ratio of the PBS fluorescence over the PSII fluorescence with betaine or glycerol added in the darkness (Supplementary Fig. S8). The effect of osmolytes on the connection between the PBS and PSII, therefore, explains the peculiar rise of NPQ with excitation of PBSs when betaine or glycerol was added under illumination. Accordingly, the degree of energetic decoupling enhanced by betaine or glycerol reflects an increased distance between the PBS and PSII. Overall, the energetic decoupling of PBSs from PSII is proposed to involve conformational change related to the connection between the PBS and PSII, which can be revealed by the reactions with stabilizing osmolytes.

NPQ is generally considered to be regulated by the transmembrane proton gradient (ΔpH) and/or the redox state of the plastoquinone pool, yet the dark recovery of the red algal NPQ is not sufficiently explained by either of the two factors. Rapid buildup of ΔpH or rapid reduction of plastoquinones with a very slow dissipation or oxidation rate, respectively, was previously proposed to be related with rapid induction of fluorescence quenching by consecutive saturating light pulses (Delphin et al., 1998; Kowalczyk et al., 2013). Based on these proposed regulatory mechanisms, fluorescence quenching is recovered due to either dissipation of ΔpH or oxidation of plastoquinols in the darkness, and the extent of the recovery is supposed to increase with the period of darkness. However, the NPQ level was enhanced by the period of dark interval (Supplementary Fig. S7A). The enhancement of NPQ over the period of darkness suggested that factors other than ΔpH and reduced plastoquinols drove fluorescence quenching, provided that the defined times of saturating light pulses induced the same extent of ΔpH and the same level of plastoquinone reduction. One possible factor that modulates the amplitude of NPQ is the surface charge on the thylakoid membrane, which has been extensively studied in isolated chloroplasts using various ionic solutions (Kaňa and Govindjee, 2016). The effect of surface charges on NPQ is compatible with the hypothesis that nigericin suppresses NPQ by influencing electrostatics of PSII and PSI (Kowalczyk et al., 2013). In this work, the effect of the cation ionophore nonactin on the NPQ kinetics supports the role of surface charges in the modulation of NPQ. The opposite enhancement and suppression effects of nonactin on the NPQ level under illumination at 300 and 10 μmol photons m^-2^ s^-1^, respectively, can be attributed to different electrochemical gradients of cations other than protons, assuming that difference in the number of ions between the lumen and the stroma is correlated to the NPQ level. Photosynthetic electron transfer promotes a rapid proton influx into the lumen followed by influx of anions to the lumen and efflux of cations other than protons to the stroma (Avenson et al., 2005). The light-acclimated amplitude of ΔpH and the corresponding anion influx and cation efflux were reported to increase over PPFD (Klughammer et al., 2013). Accordingly, the electrochemical potential of cations other than protons in the stroma relative to the lumen becomes more positive under high illumination than under low illumination. Therefore, it is likely that nonactin induces cation influx to the lumen under high illumination and cation efflux to the stroma under low illumination, leading to the opposite effects on the NPQ level. Furthermore, the membrane potential (ΔΨ) caused by difference in the concentrations of all ions across the membrane likely accounts for the residual NPQ in the presence of nigericin under illumination at 300 μmol photons m^-2^, as nigericin dissipates ΔpH and maintains ΔΨ. This residual NPQ is further abolished by triclosan that dissipates both ΔpH and ΔΨ. The involvement of the surface charge in the modulation of the NPQ level in the darkness is also supported by that the dark relaxation of NPQ is facilitated by nigericin and/or nonactin (Fig. 7A-B). Taken together, these observations indicate that fluorescence quenching is modulated by electrochemical gradient of ions across the membrane most likely through influencing the surface charge on the thylakoid membrane in red algae.

The energetic decoupling of PBSs from PSII likely involves a regulatory mechanism distinct from the other quenching processes since the former is specifically enhanced by CCCP. The distinctness of the regulatory mechanism can also be inferred from the kinetics of the energy decoupling. Whereas the energy decoupling stayed at a nearly steady degree after the rapid induction by light, the changes in the other fluorescence quenching processes, as judged by the NPQ level, were dynamic depending on the illumination period. The degree of the energy decoupling was slightly enhanced by high illumination at 300 μmol photons m^-2^ s^-1^, while the light-acclimated NPQ level at 6 min of illumination drastically varied over PPFD. Based on the mechanistic interpretation of the energy decoupling discussed before, the enhancement of the energy decoupling by CCCP reflects the increase in the distance between the PBS and PSII and may thus involve conformational changes of either or both of the PBS and PSII. CCCP was reported to influence the electron transfer in PSII by dampening the period four oscillation of oxygen evolution and prolonging the lifetime of Q_A_^-^ (de Wijn and van Gorkom, 2001). Furthermore, light-induced electron transfer kinetics were reported to be associated with conformational changes of PSII (Kulik et al., 2020). Therefore, CCCP may affect the protein conformation of PSII, leading to a loose connection between the PBS and PSII. The influence of CCCP on the degree of energetic decoupling is prominent under high illumination at 300 μmol photons m^-2^ s^-1^, which can be attributed to a presumably larger conformation change of PSII related to electron transfer than that under low illumination. Furthermore, the counteracting effect of DCMU on the CCCP-enhanced energetic decoupling is likely mediated by suppression of the conformation changes related to the electron transfer within PSII. Taken together, the effect of CCCP on NPQ supports that the modulation of the energetic decoupling from PBSs to PSII is associated with conformational changes associated to the connection between the PBS and PSII which are responsive to light and tuned by light intensity.

## CONCLUSION

By using light preferentially absorbed by PBSs or by photosystems, the fluorescence quenching processes associated with the PBSs was distinguishable from those associated with PSII. As a result, the energetic decoupling of PBSs from PSII was identified as being involved in NPQ along with the previously identified intrinsic PSII quenching in *C. merolae*. The energetic decoupling of PBSs from PSII was mechanistically distinguished from other fluorescence quenching processes by stabilizing osmolytes and likely involved conformational changes related to the connection between the PBS and PSII. This finding provides a novel mechanistic view of the interaction between the PBS and PSII involved in photosynthetic regulation of red alga. The effects of various ionophores on the NPQ kinetics suggest that NPQ is modulated by the surface charge on thylakoid membranes in addition to the transmembrane gradient of protons. By contrast, the redox state of plastoquinone pool did not apparently modulate the NPQ level, as determined by similar suppression of NPQ by DCMU and DBMIB. The multiple NPQ processes distinguished by their kinetics and specific chemicals further the understanding of the mechanism underlying modulation of red algal NPQ that is unique and different from those identified in plants, green algae, and cyanobacteria.

## MATERIALS AND METHODS

### Strains and culture conditions

The red algal strain *Cyanidioschyzon merolae* 10D (NIES-3377) was obtained from the Microbial Culture Collection at the National Institute for Environmental Studies (http://mcc.nies.go.jp; Tsukuba, Japan) and grown in a MA liquid medium (pH adjusted to 2.0 with H_2_SO_4_; Minoda et al., 2004). The cells were cultured on a rotary shaker (125 rpm) at 40°C under continuous cool-white fluorescent light (20 µmol photons m^−2^ s^−1^). The cells were harvested in the logarithmic phase of growth for all the following experiments.

### Fluorescence spectrometry

Fluorescence-based kinetics were measured using a MultispeQ v2.0 spectrophotometer (PhotosynQ). Cells were centrifuged at 3,000×g for 2 min and resuspended in a fresh MA liquid medium with the chlorophyll concentration adjusted to 15 μg mL^−1^ unless otherwise indicated. The resuspended cells were incubated on a rotary shaker (125 rpm) in darkness at 34°C for at least 2 h. Before each measurement, 1.5 mL of cells were placed in a 1×1-cm square cuvette clamped between the light guides and incubated in darkness at the indicated temperature for 20 min. During the measurement, cells were constantly stirred at 250 rpm to prevent sedimentation.

NPQ kinetics were measured using a red light-emitting diode (LED) as actinic light source. Fluorescence levels were recorded using an amber LED (30 ns × 400 μmol photons m^−2^ s^−1^) as detecting light source unless otherwise indicated. Signals in the cuvette without cells added arose from detecting light and were recorded for baseline correction of the fluorescence level. A saturating light pulse (160 ms × 8000 μmol photons m^−2^ s^−1^, red LED) was applied to measure the maximum fluorescence level. NPQ is equivalent to (F_M_ / F_M_’) − 1, where F_M_ is the maximum fluorescence level in the dark-acclimated condition and Fm’ the maximum fluorescence level after actinic light illumination. PSII operating efficiency, Φ_PSII_, is calculated from (F_M_’ – F_S_’) / F_M_’, where F_S_’ is the steady-state fluorescence level before application of saturating light (Baker, 2008). Fluorescence kinetics using detecting lights of different colors were measured according to the manufacturer’s guideline (https://help.photosynq.com/protocols/pulses.html#pulse-sets). Fluorescence levels were sequentially and repeatedly recorded using green, amber, and blue detecting light pulses (30 ns × 500 μmol photons m^−2^ s^−1^ each pulse).

The functional antenna size of PSII with excitation of PSII or PB was estimated from fluorescence induction curves using blue or amber actinic light, respectively. For each color of actinic light, light intensity was set to 200 μmol photons m^−2^ s^−1^, and the total period of actinic light was 400 ms. Fluorescence levels were recorded using a green LED (30 ns × 400 μmol photons m^−2^ s^−1^) as detecting light source. To obtain a good signal to noise ratio of the fluorescence level, cells were resuspended in a fresh MA liquid medium with the chlorophyll concentration adjusted to 30 μg mL^−1^. To estimate the functional antenna size at different periods of light and/or darkness treatment, 3-(3,4-dichlorophenyl)-1,1-dimethylurea (DCMU, 20 μM) was added under the dark-acclimated condition or immediately after the last saturating light pulse was applied, and fluorescence induction curves were recorded 20 s after the addition of DCMU. The values of fluorescence induction curve were normalized to the fluorescence level before actinic light as 0 and to the maximum fluorescence level of the curve as 1. The functional antenna size was calculated as the reciprocal of the proportion of the area above the normalized fluorescence induction curve.

The stock solution of osmolytes, ionophores and other inhibitors was prepared in the MA liquid medium (betaine, glycerol), ethanol (triclosan, DCMU, DBMIB), or ethanol:dimethyl sulfoxide 1:1 (v/v) (nigericin, nonactin, CCCP). The effective concentration of ionophores and inhibitors was determined by the maximum effect on the NPQ kinetics and was 5 μM for nigericin, CCCP, DBMIB, 10 μM for nonactin, and 20 μM for triclosan and DCMU.

### 77K fluorescence emission spectroscopy

Fluorescence emission spectra were recorded in an adapted stainless sample chamber using a FLAME-S spectrometer (Ocean Optics). Laser beam with a wavelength of either 440 nm (preferentially exciting PSI and PSII; CivilLaser LSR440NL-1W) or 589 nm (preferentially exciting PBS; CivilLaser LSR589H-100) was generated as an excitation light source. Light exiting the laser apparatus was collimated and passed through an optic fiber of bifurcated fiber bundle (ThorLabs BFY1000HS02) to the sample chamber. Light emitted from the sample chamber was collected by the other optical fiber of the fiber bundle and passed through a long pass emission filter with a cut-off wavelength of either 465 nm (for 440-nm excitation light; Chroma AT465lp) or 610 nm (for 589-nm excitation light; Chroma AT610lp) to the spectrometer. To capture the fluorescence emission spectrum at different periods of light and/or darkness treatment, 4 μL of cells were collected and immediately mixed with 26 μL of exogenous fluorophore on ice. The final chlorophyll concentration in the mixed solution was 2 μg mL^−1^; the final concentration of exogenous fluorophore for normalization was 0.8 μM of Rhodamine 6G for 440-nm excitation light and 13.52 μM of acid-coated water soluble 900 nm PbS/CdS quantum dots (PBS900-WS-AC, NanoOptical Materials) for 589-nm excitation light. The mixed solution was then placed in the ice-cold sample chamber and covered by a circular cover glass (10 mm in diameter). The sample chamber was connected to the fiber bundle and equilibrated with liquid nitrogen. During the procedures from sample collection until placement of the sample chamber in liquid nitrogen, the sample was constantly kept on ice under dim light (<1 μmol photons m^−2^ s^−1^). The fluorescence spectrum was corrected by subtraction with the baseline levels due to light scattering and then by normalization to the maximum fluorescence amplitude of exogenous fluorophore at a peak wavelength (542 nm for Rhodamine 6G and 890 nm for PbS/CdS quantum dots) as 1.

To quantify the fluorescence amplitude of different components, the emission spectra subtracted by the normalized emission spectrum of exogenous fluorophores were fitted with Gaussian curves using OriginPro 2018 (OriginLab), and the fluorescence emission components were assigned based on the peak wavelength of the Gaussian curve according to the literature (Scott et al., 2006; Lamb et al., 2018). As fluorescence emission components of CP43 (F685) and CP47 (F695) with excitation wavelength at 440 nm were difficult to be deconvoluted and involved large deviation of full width at half maximum in few cases, the boundary conditions were set for F685 (between 5 and 15 nm) and F695 (between 5 and 10 nm). No boundaries were set for the other fitting parameters of the Gaussian curve.

## Acknowledgements

We thank Shao-Lun Liu at Tunghai University, Taiwan, for his kind gift of the *C. merolae* 10D strain. This work was supported by Ministry of Science and Technology, Taiwan (MOST 108-2311-B-110-001-, 109-2311-B-110-002-, 110-2311-B-110-002-MY3) and by the Higher Education Sprout Project, Taiwan.

## Reference

Avenson TJ, Kanazawa A, Cruz JA, Takizawa K, Ettinger WE, Kramer DM (2005) Integrating the proton circuit into photosynthesis: progress and challenges. Plant Cell Environ 28: 97–109

Baker NR (2008) Chlorophyll fluorescence: a probe of photosynthesis in vivo. Annu Rev Plant Biol 59: 89–113

Barbosa O, Ortiz C, Berenguer-Murcia Á, Torres R, Rodrigues RC, Fernandez-Lafuente R (2014) Glutaraldehyde in bio-catalysts design: a useful crosslinker and a versatile tool in enzyme immobilization. RSC Advances 4: 1583–1600

Belgio E, Kapitonova E, Chmeliov J, Duffy CDP, Ungerer P, Valkunas L, Ruban AV (2014) Economic photoprotection in photosystem II that retains a complete light-harvesting system with slow energy traps. Nat Commun 5: 4433

Bhatti AF, Choubeh RR, Kirilovsky D, Wientjes E, van Amerongen H (2020) State transitions in cyanobacteria studied with picosecond fluorescence at room temperature. Biochim Biophys Acta - Bioenerg 1861: 148255

Bhatti AF, Kirilovsky D, van Amerongen H, Wientjes E (2021) State transitions and photosystems spatially resolved in individual cells of the cyanobacterium *Synechococcus elongatus*. Plant Physiol 186: 569–580

Busch A, Nield J, Hippler M (2010) The composition and structure of photosystem I-associated antenna from *Cyanidioschyzon merolae*. Plant J 62: 886–897

Calzadilla PI, Kirilovsky D (2020) Revisiting cyanobacterial state transitions. Photochem Photobiol Sci

Calzadilla PI, Zhan J, Sétif P, Lemaire C, Solymosi D, Battchikova N, Wang Q, Kirilovsky D (2019) The cytochrome *b*_6_*f* complex Is not involved in cyanobacterial state transitions. Plant Cell 31: 911–931

Chang L, Liu X, Li Y, Liu C-C, Yang F, Zhao J, Sui S-F (2015) Structural organization of an intact phycobilisome and its association with photosystem II. Cell Res 25: 726–737

de Wijn R, van Gorkom HJ (2001) Kinetics of electron transfer from Q_A_ to Q_B_ in photosystem II. Biochemistry 40: 11912–11922

Delphin E, Duval J-C, Etienne A-L, Kirilovsky D (1996) State transitions or ΔpH-dependent quenching of photosystem II fluorescence in red algae. Biochemistry 35: 9435–9445

Delphin E, Duval J-C, Etienne A-L, Kirilovsky D (1998) ΔpH-dependent photosystem II fluorescence quenching induced by saturating, multiturnover pulses in red algae. Plant Physiol 118: 103–113

Derks A, Schaven K, Bruce D (2015) Diverse mechanisms for photoprotection in photosynthesis. Dynamic regulation of photosystem II excitation in response to rapid environmental change. Biochim Biophys Acta - Bioenerg 1847: 468–485

Gardian Z, Bumba L, Schrofel A, Herbstova M, Nebesarova J, Vacha F (2007) Organisation of photosystem I and photosystem II in red alga *Cyanidium caldarium*: encounter of cyanobacterial and higher plant concepts. Biochim Biophys Acta 1767: 725–731

Haniewicz P, Abram M, Nosek L, Kirkpatrick J, El-Mohsnawy E, Olmos JDJ, Kouřil R, Kargul JM (2018) Molecular mechanisms of photoadaptation of photosystem I supercomplex from an evolutionary cyanobacterial/algal intermediate. Plant Physiol 176: 1433–1451

Homann PH (1971) Actions of carbonylcyanide m-chlorophenylhydrazone on electron transport and fluorescence of isolated chloroplasts. Biochim Biophys Acta - Bioenerg 245: 129–143

Kaňa R, Govindjee (2016) Role of ions in the regulation of light-harvesting. Front Plant Sci 7: 1849–1849

Kaňa R, Kotabová E, Lukeš M, Papáček Š, Matonoha C, Liu L-N, Prášil O, Mullineaux CW (2014) Phycobilisome mobility and its role in the regulation of light harvesting in red algae. Plant Physiol 165: 1618–1631

Klughammer C, Siebke K, Schreiber U (2013) Continuous ECS-indicated recording of the proton-motive charge flux in leaves. Photosynthesis Res 117: 471–487

Koblížek M, Komenda J, Masojídek J (1998) State transitions in the cyanobacterium *Synechococcus* PCC 7942. Mobile antenna or spillover? In G Garab, ed, Photosynthesis: Mechanisms and Effects: Volume I–V: Proceedings of the XIth International Congress on Photosynthesis, Budapest, Hungary, August 17–22, 1998. Springer Netherlands, Dordrecht, pp 213-216

Kowalczyk N, Rappaport F, Boyen C, Wollman FA, Collen J, Joliot P (2013) Photosynthesis in *Chondrus crispus*: the contribution of energy spill-over in the regulation of excitonic flux. Biochim Biophys Acta 1827: 834–842

Krasilnikov PM, Zlenko DV, Stadnichuk IN (2020) Rates and pathways of energy migration from the phycobilisome to the photosystem II and to the orange carotenoid protein in cyanobacteria. FEBS Lett 594: 1145–1154

Krupnik T, Kotabova E, van Bezouwen LS, Mazur R, Garstka M, Nixon PJ, Barber J, Kana R, Boekema EJ, Kargul J (2013) A reaction center-dependent photoprotection mechanism in a highly robust photosystem II from an extremophilic red alga, *Cyanidioschyzon merolae*. J Biol Chem 288: 23529–23542

Kulik N, Kutý M, Řeha D (2020) The study of conformational changes in photosystem II during a charge separation. J Mol Model 26: 75

Lamb JJ, Røkke G, Hohmann-Marriott MF (2018) Chlorophyll fluorescence emission spectroscopy of oxygenic organisms at 77 K. Photosynthetica 56: 105–124

Ley AC, Butler WL (1977) Energy transfer from photosystem II to photosystem I in *Porphyridium cruentum*. Biochim Biophys Acta - Bioenerg 462: 290–294

Li X-P, Björkman O, Shih C, Grossman AR, Rosenquist M, Jansson S, Niyogi KK (2000) A pigment-binding protein essential for regulation of photosynthetic light harvesting. Nature 403: 391–395

Li Y, Zhang J, Xie J, Zhao J, Jiang L (2001) Temperature-induced decoupling of phycobilisomes from reaction centers. Biochim Biophys Acta - Bioenerg 1504: 229–234

Liu L-N, Aartsma TJ, Thomas J-C, Zhou B-C, Zhang Y-Z (2009) FRAP analysis on red alga reveals the fluorescence recovery is ascribed to intrinsic photoprocesses of phycobilisomes than large-scale diffusion. PLoS One 4: e5295

Liu L-N, Elmalk AT, Aartsma TJ, Thomas J-C, Lamers GEM, Zhou B-C, Zhang Y-Z (2008) Light-induced energetic decoupling as a mechanism for phycobilisome-related energy dissipation in red algae: a single molecule study. PLoS One 3: e3134

McConnell MD, Koop R, Vasil’ev S, Bruce D (2002) Regulation of the distribution of chlorophyll and phycobilin-absorbed excitation energy in cyanobacteria. A structure-based model for the light state transition. Plant Physiol 130: 1201–1212

Mehra R, Dehury B, Kepp KP (2020) Cryo-temperature effects on membrane protein structure and dynamics. Phys Chem Chem Phys 22: 5427–5438

Minoda A, Sakagami R, Yagisawa F, Kuroiwa T, Tanaka K (2004) Improvement of culture conditions and evidence for nuclear transformation by homologous recombination in a red alga, *Cyanidioschyzon merolae* 10D. Plant Cell Physiol 45: 667–671

Murata N (1969) Control of excitation transfer in photosynthesis I. Light-induced change of chlorophyll a fluoresence in *Porphyridium cruentum*. Biochim Biophys Acta - Bioenerg 172: 242–251

Nicol L, Nawrocki WJ, Croce R (2019) Disentangling the sites of non-photochemical quenching in vascular plants. Nat Plants 5: 1177–1183

Papageorgiou GC, Govindjee, Govindjee R, Mimuro M, Stamatakis K, Alygizaki-Zorba A, Murata N (1999) Light-induced and osmotically-induced changes in chlorophyll a fluorescence in two *Synechocystis* sp. PCC 6803 strains that differ in membrane lipid unsaturation. Photosynthesis Res 59: 125–136

Peers G, Truong TB, Ostendorf E, Busch A, Elrad D, Grossman AR, Hippler M, Niyogi KK (2009) An ancient light-harvesting protein is critical for the regulation of algal photosynthesis. Nature 462: 518–521

Rösgen J, Pettitt BM, Bolen DW (2005) Protein Folding, Stability, and Solvation Structure in Osmolyte Solutions. Biophys J 89: 2988–2997

Ruban AV, Johnson MP (2009) Dynamics of higher plant photosystem cross-section associated with state transitions. Photosynthesis Res 99: 173–183

Scott M, McCollum C, Vasil’ev S, Crozier C, Espie GS, Krol M, Huner NPA, Bruce D (2006) Mechanism of the down regulation of photosynthesis by blue light in the cyanobacterium *Synechocystis* sp. PCC 6803. Biochemistry 45: 8952–8958

Sonoike K (2011) Photoinhibition of photosystem I. Physiol Plant 142: 56–64

Tian L, Nawrocki WJ, Liu X, Polukhina I, van Stokkum IHM, Croce R (2019) pH dependence, kinetics and light-harvesting regulation of nonphotochemical quenching in *Chlamydomonas*. Proc Natl Acad Sci USA 116: 8320–8325

Tibiletti T, Auroy P, Peltier G, Caffarri S (2016) *Chlamydomonas reinhardtii* PsbS protein is functional and accumulates rapidly and transiently under high light. Plant Physiol 171: 2717–2730

Ueno Y, Aikawa S, Niwa K, Abe T, Murakami A, Kondo A, Akimoto S (2017) Variety in excitation energy transfer processes from phycobilisomes to photosystems I and II. Photosynthesis Res: 1–9

Wilson A, Ajlani G, Verbavatz J-M, Vass I, Kerfeld CA, Kirilovsky D (2006) A soluble carotenoid protein involved in phycobilisome-related energy dissipation in cyanobacteria. Plant Cell 18: 992–1007

Wollman F-A (2001) State transitions reveal the dynamics and flexibility of the photosynthetic apparatus. EMBO J 20: 3623–3630

